# Circulating Clonally Expanded T Cells Reflect Functions of Tumor Infiltrating T Cells

**DOI:** 10.1101/2020.09.30.321240

**Authors:** Liliana Lucca, Pierre-Paul Axisa, Benjamin Lu, Brian Harnett, Shlomit Jessel, Le Zhang, Khadir Raddassi, Lin Zhang, Kelly Olino, James Clune, Meromit Singer, Harriet M. Kluger, David A. Hafler

## Abstract

Understanding the relationship between tumor and peripheral immune environments could allow longitudinal immune monitoring in cancer. Here, we examined whether T cells that share the same TCRαβ and are found in both tumor and blood can be interrogated to gain insight into the ongoing tumor T cell response. Paired transcriptome and TCRαβ repertoire of circulating and tumor-infiltrating T cells were analyzed from matched tumor and blood from patients with metastatic melanoma at the single cell level. We found that in circulating T cells matching clonally expanded tumor-infiltrating T cells (circulating TILs), gene signatures of effector functions, but not terminal exhaustion, reflect those observed in the tumor. In contrast, features of exhaustion are displayed predominantly by T cells present only in tumor. Finally, genes associated with a high degree of blood-tumor TCR sharing were overexpressed in tumor tissue after immunotherapy. These data demonstrate that circulating TILs, identified by TCRs shared with T cells in tumors, have unique transcriptional expression patterns that may have utility for the interrogation of T cell function in cancer immunotherapy.

**Summary:** Combining transcriptomic and TCRαβ repertoire analysis of circulating and tumor-infiltrating CD8 T cells from patients with metastatic melanoma, we identify a blood-based population with effector properties that reflect those of clonally related tumor-resident T cells.

## Introduction

Tumor infiltration by T cells, particularly CD8^+^ cells, is a significant prognostic factor in cancer (reviewed in Barnes and Amir, 2017). Cytotoxic T cells, characterized by lytic granule components and NK-associated receptors, play a pivotal role in tumor rejection by directly destroying tumor cells and associated structures in an antigen-specific fashion (Breart et al., 2008; Schietinger et al., 2013). In contrast, exhausted T cells are defined by sequential loss of effector functions, limited proliferative potential (Blank et al., 2019; Moskophidis et al., 1993) and increased, stable expression of co-inhibitory receptors (reviewed in Blank et al., 2019; van der Leun et al., 2020). Inhibition of co-inhibitory signals has proven highly successful in the treatment of solid malignancies (Ribas and Wolchok, 2018). Understanding the relationship between tumor and peripheral immune environments could allow longitudinal immune monitoring in situations in which tumor biopsies are unfeasible (Cohen and Buchbinder, 2019). Moreover, serial sampling for predicting and monitoring response to therapy is often necessary when weighing risk/benefit ratios for continuation of treatment.

Here, we analyzed paired transcriptome and TCRαβ repertoire of circulating and tumor-infiltrating T cells from matched tumor and blood samples from patients with metastatic melanoma at the single cell level. We tested the hypothesis that circulating sister clones of clonally expanded TILs, i.e. clones with matching TCR sequences, can be interrogated to gain insight into the ongoing T cell response in the tumor. We found that the degree of cytotoxicity of the tumor infiltrate, but not exhaustion, can be inferred from circulating TILs. The most pronounced features of exhaustion are consistently displayed by tumor-exclusive clonal T cells. Thus, the degree of TCR repertoire sharing across tissues predicts the T cell phenotype, and genes associated with a high degree of blood-tumor TCR sharing are overexpressed by tissue-infiltrating T cells after successful immunotherapy.

## Results and Discussion

### Detection of clonally-expanded TILs in the circulation of melanoma patients

We analyzed paired transcriptomic and TCRαβ libraries from a total of 123,868 T cells derived from matched metastatic tumors and blood from 11 stage IV melanoma patients selecting subjects with diverse treatment history (Table 1). A single TCRαβ clonotype (see Methods) could be detected in 66.1% (48.5% – 76.6%) of the cells, and only these cells were retained for subsequent analyses. A reduction in the diversity of immune repertoires signals an ongoing immune response. We computed four estimates of repertoire diversity and compared tumor and blood repertoires across patients and observed a significant reduction in the richness of clonotype species in the tumor as calculated by the Chao (Chao et al., 1992) and ACE indexes (Laydon et al., 2014) (Fig 1A). To identify patterns of similarity, we performed hierarchical clustering based on clonal size distribution and observed an overall separation of blood and tumor samples with two tumor samples, Mel UT-2 and Mel T-2, clustering closer to blood samples (Fig. 1B). These two samples consistently ranked lowest when measuring the proportion of the repertoire occupied by the top 10 most frequent clonotypes (Fig 1C). Of note, Mel UT-2 was a mucosal melanoma and therefore sun-shielded (Table 1), consistent with the correlation between UV radiation exposure and immune infiltration (Wang et al., 2017). These analyses confirmed previous results showing restricted TCR repertoires in melanoma, indicative of ongoing antigen-specific responses (Kalaora et al., 2018). We then classified individual TIL clonotypes into clonally expanded or not, based on the proportion of the repertoire occupied by each clonotype. Because of significant variability in the number of analyzed TILs across patients (1,127 – 5,404), for this classification we established individual percentage-based thresholds (Fig. S1). We next examined whether sister clones of expanded TILs could be detected in the blood based on exact clonotype matching. Comparing the proportion of tumor and blood repertoires occupied by the 20 most expanded TIL clonotypes highlights the existence of both shared and tumor exclusive large clones (Fig 1D). Based on the largely mutually exclusive expression of the *CD8A* and *CD4* genes, we found that both tumor-resident and circulating expanded TILs were disproportionally CD8 (Fig 1E-G). Finally, we observed a positive correlation between the frequency of circulating TILs in blood, (average 7.5% ± 10.4%, Fig 1H) and the duration of metastatic disease (Table 1 and Fig 1I).

**Fig. 1.**
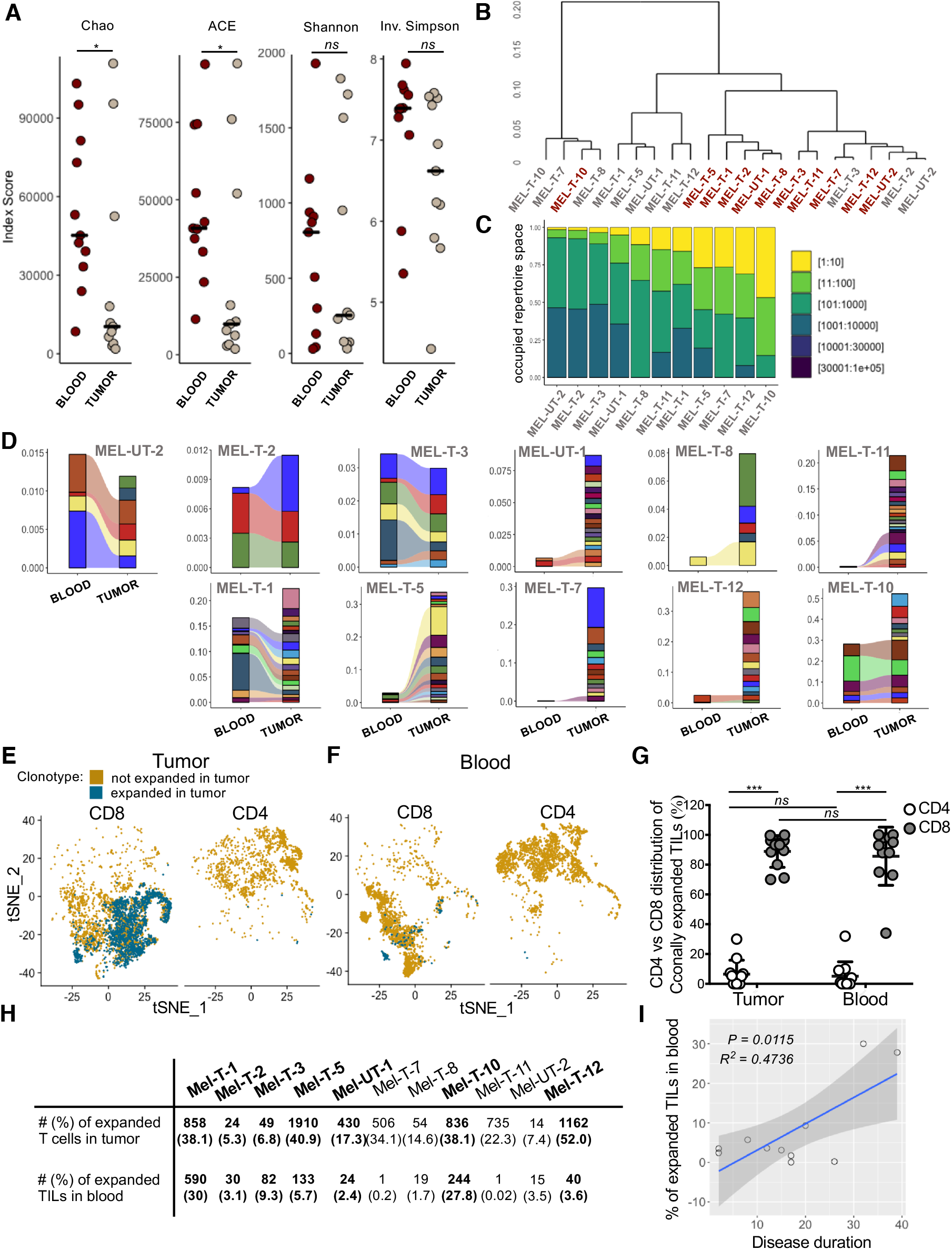
Detection of clonally-expanded TILs in the circulation of melanoma patients. **A**, dot plots displaying indexes of diversity and species richness for tumor and blood TCRαβ repertoires of individual patients. Horizontal bars indicate medians. Statistical significance was determined using Wilcoxon rank-sum test. **B**, hierarchical clustering of tumor and blood TCRαβ repertoires based on clonal size distribution. Red = blood, Grey = tumor. **C**, bar graph displaying the proportion of TCRαβ repertoire space occupied by the top n clonotypes (bins in legend) for the tumor samples of individual patients. D, alluvial plots displaying the frequency of the top 20 most expanded TIL clonotypes in blood and tumor for individual patients. E-F, tSNE plot of TILs (**B**) and circulating T cells (**C**) divided by expression of CD8A (normalized counts > 1) and CD4 (normalized counts > 1) for one representative patient. T cells expressing TCRs expanded in the tumor are highlighted in blue. **D**, table displaying the absolute number and frequency of tumor-resident and blood-derived CD8 T cells expressing a TCR expanded in the tumor for each patient. Bold indicates patients in which the absolute number of blood-derived T cells with a TCR expanded in the tumor (circulating TILs) was deemed large enough for subsequent analyses. **E**, percentage of CD4+ (white) and CD8+ (grey) T cells among clonally expanded T cells in tumor and circulating TILs. Each data point represents the average for one patient. Data represent mean +/- standard deviation. Statistical significance was determined using t test with Bonferroni-Dunn correction for multiple comparisons. **F** scatter plots showing the relationship of the frequency of circulating TILs and disease duration. Adjusted R-squared and F test p value are shown.

**Table 1.** Demographics and clinical features of patients included in the study. All patients are stage IV melanoma (Gershenwald et al., 2017). *Tumor burden* is defined based on a lesion volume threshold of 2 cm^3^. *Disease duration* corresponds to the time between the diagnosis of metastatic disease and sample collection. *Response* is defined according to the RECIST guidelines (v1.1, (Eisenhauer et al., 2009). N.A. = not applicable, P.D. = progressive disease, P.R. = partial response. <u>*Tumor burden* was calculated using the formula (4/3)π r3, where r is the maximum radius of each metastasis, except for three lesions for which the formula for the volume of an ellipsoid was deemed more appropriate (V=4/3π*L/2*H/2*W/2)</u>.

| Patient I.D. | Sex | Age | Lesion location | BRAF/NRAS/c-Kit mutation | Sun exposure | Tumor burden | Disease duration (mo) |  | ICB history | Most recent ICB infusion | Response |
| --- | --- | --- | --- | --- | --- | --- | --- | --- | --- | --- | --- |
| Mel-T-1 | F | 62 | Lymph node | WT | Exposed | Large | 32 |  | Treated | Pembrolizumab | P.D. |
| Mel-T-2 | M | 40 | Lymph node | WT | Exposed | Large | 15 (IIIC) |  | Treated | APX005M + Cabiralizumab + | P.D. |
| Mel-T-3 | M | 71 | Lymph node | NRAS | Exposed | Small | 20 (IIIB) |  | Treated | Nivolumab | P.D. |
| Mel-T-5 | F | 44 | Lymph node | NRAS | Exposed | Large | 8 |  | Treated | Pembrolizumab | P.D. |
| Mel-T-6 | M | 57 | Lung | NRAS | Exposed | Small | 29 |  | Treated | Nivolumab + Ipilimumab | P.R. |
| Mel-UT-1 | F | 79 | Lymph node | BRAF | Exposed | Small | 2 (IIIB) |  | Naïve | N.A. | N.A. |
| Mel-T-7 | F | 40 | Gall bladder | BRAF | Exposed | Large | 26 |  | Treated | APX005M + Cabiralizumab + | P.D. |
| Mel-T-8 | M | 68 | Small bowel | BRAF | Exposed | Large | 17 |  | Treated | Pembrolizumab | P.D. |
| Mel-T-9 | M | 43 | Lymph node | BRAF | Exposed | Small | 22 |  | Treated | Nivolumab + Ipilimumab | P.R. |
| Mel-T-10 | M | 61 | Soft tissue | BRAF | Exposed | Large | 39 |  | Treated | APX005M + Cabiralizumab + | P.D. |
| Mel-T-11 | M | 36 | Lymph node | BRAF | Exposed | Small | 17 |  | Treated | Nivolumab + Ipilimumab | P.D. |
| Mel-UT-2 | M | 71 | Lymph node | WT | Shielded (mucosal) | Large | 2 (IIID) |  | Naïve | N.A. | N.A. |
| Mel-T-12 | M | 34 | Skin metastasis | BRAF | Exposed | Large | 12 |  | Treated | Nivolumab + Ipilimumab | P.R. |

### Cytotoxic profile of circulating TILs

We then examined whether clonally expanded T cells in the tumor displayed specific functional traits by analyzing transcriptional profiles of individual CD8 T cells from all patients by grouping similar cells into clusters. We identified seventeen clusters (Fig 2A), and clonally expanded TILs largely map onto clusters overrepresented in the tumor (Fig 2B). We further annotated each cluster by identifying cluster-specific gene markers (Table S1) and comparing these markers with gene sets derived from previous studies of the immune infiltrate in melanoma at the single cell level (Li et al., 2019; Sade-Feldman et al., 2018; Tirosh et al., 2016). A large subgroup of blood-dominated clusters (6, 13, 16, 8) corresponded to naïve and central memory CD8 T cells, defined by the high expression of *CCR7* and *IL7R*, overlapping with published signatures of central memory (Fig 2C-D). Clusters 2 and 12 display both signatures of central and effector memory, while clusters 3, 9 and 14 express high levels of cytotoxic molecules and cluster together with previously reported cytotoxic populations (Fig 2C-D). Tumor-dominated clusters 0, 5, 10, 15 were enriched in signatures of exhaustion and terminal differentiation and are indeed marked by expression of *PDCD1* (Fig 2C-D). Clusters 1, and 11 and presented an intermediate profile, with both signatures of exhaustion and cytotoxicity, and higher expression of *GZMK* than *GZMB* (Zhang et al., 2019) (Fig 2C-D), possibly representing pre-dysfunctional T cells (Li et al., 2018). We also identified two clusters of actively proliferating cells (4 and 7), expressing high levels of *MKI67* and predominantly containing tumor-derived, clonally expanded T cells.

**Fig. 2.**
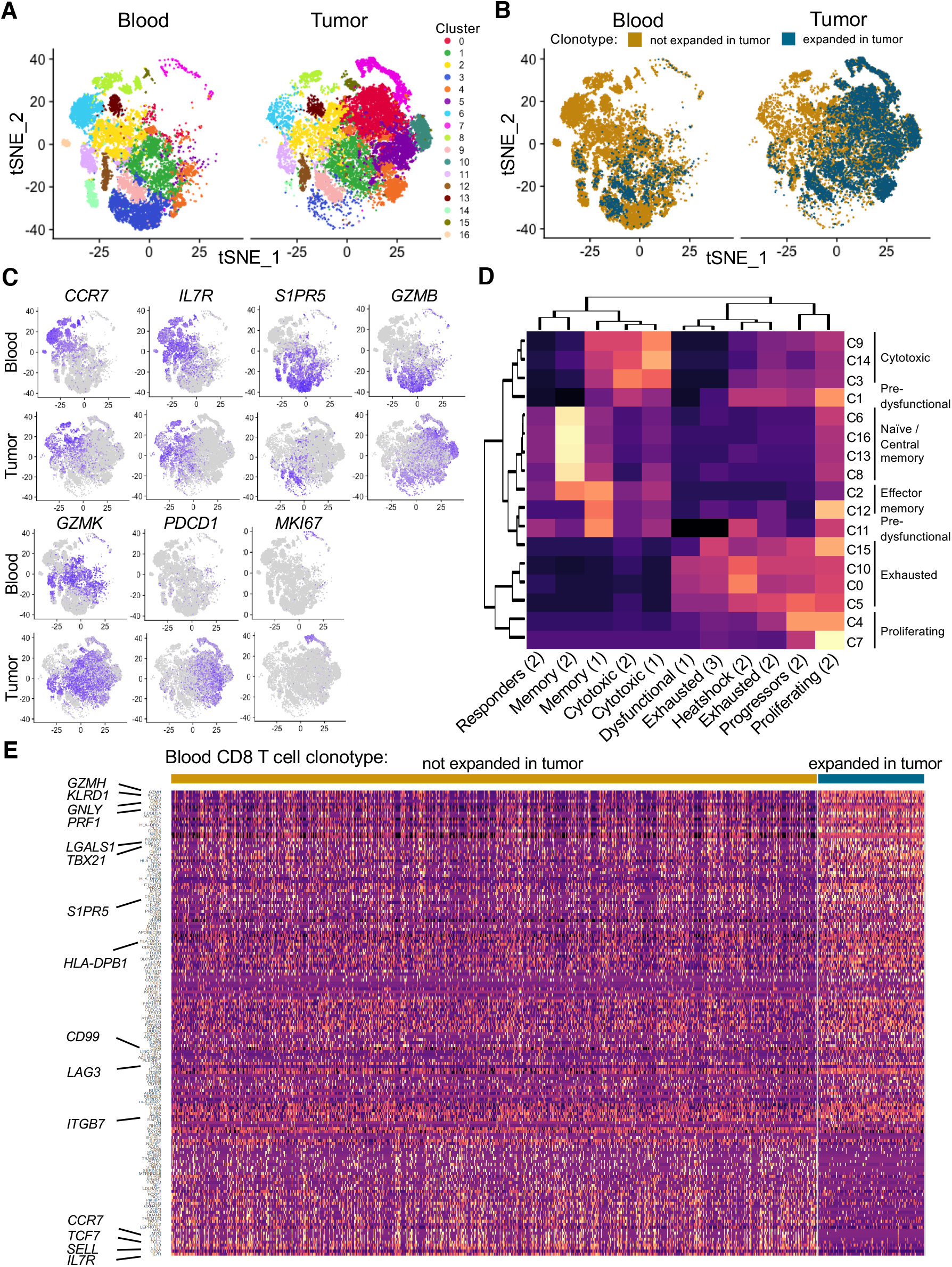
Cytotoxic profile of circulating TILs. **A**, tSNE plot of CD8 T cells from 11 patients, split by tissue of origin and coloured by cluster. Clustering was obtained with a SNN modularity optimization-based algorithm (k parameter = 5, resolution = 0.5). **B**, tSNE plot of CD8 T cells from 11 patients, split by tissue of origin and coloured by expression of a TCR expanded in the tumor (blue, threshold for expansion as in Supplementary Fig 1), or not (gold). **C**, tSNE plots coloured by scaled expression of indicated genes. **D**, clustered heatmap diplaying the degree of overlap between lists of cluster markers (rows) and gene sets derived from published studies of melanoma TIL transcriptome (1 = Li et al. 2019, 2 = Sade-Feldman et al. 2018, 3 = Tirosh et al. 2016). **E**, heatmap of differentially expressed genes between circulating TILs and blood T cells with a TCR not expanded in the tumor with a delta % of expressing cells between the two groups of at least +/-10%. Rows represent genes ordered by delta % of expressing cells. Columns represent individual cells grouped by identity class as indicated on the horizontal bars. Values are scaled and mean-centered log2-transformed gene counts. For visualization purposes, the dataset was randomly down sampled to include 1000 cells. Values are scaled and mean-centered log2-transformed gene counts.

In the blood, circulating TILs map to both cytotoxic and pre-dysfunctional clusters. To further define transcriptional traits associated with this population, we compared transcription profiles of circulating TILs and other circulating T cells. We detected higher expression of multiple cytotoxic molecules such as granzymes, perforin and NK-associated receptors, and also markers of tissue residence (*LGALS1*), migration (*CD99*) and tissue homing *(ITGB7*). Finally, *LAG3*, but not other coinhibitory receptors, was also significantly upregulated by circulating TILs (Fig 2E).

### Tumor signature of circulating TILs

We then examined whether circulating TILs are enriched for features of tumor resident, clonally expanded TILs. We focused on seven patients where a minimum of 20 circulating TILs could be detected (Fig 1H). For each patient, we defined expanded TIL gene sets consisting of genes differentially expressed between clonally expanded TILs and tumor-resident T cells with a unique TCR (singletons, Fig 3A). Similarly, we compared clonally expanded, blood-derived T cells (based on thresholds shown in Fig. S2) and blood-derived singletons (Fig 3A). To obtain a list of genes specifically associated with clonal expansion in the tumor, we incorporated a term for the interaction between clonal expansion and tissue into the linear model used for differential expression detection (Fig 3B). This analysis yielded two types of gene sets: those containing all genes relevant to clonal expansion in the tumor, including hallmarks of general activation (expanded TIL gene sets), and those containing only genes found exclusively when T cells expand in the tumor (tumor-specific gene sets, Table S3). We then calculated the overlap between expanded TIL gene sets from individual patients. The expanded TIL gene sets of Met-T-1 and Mel-T-2, the two patients with BRAF/NRAS/c-kit wild-type tumors, presented a high degree of overlap and shared multiple markers of cytotoxicity. A high degree of similarity was also observed for subjects Mel-T-5 and Mel-T-12, enriched for signatures of exhaustion (Fig 3C, Table S3).

**Fig. 3.**
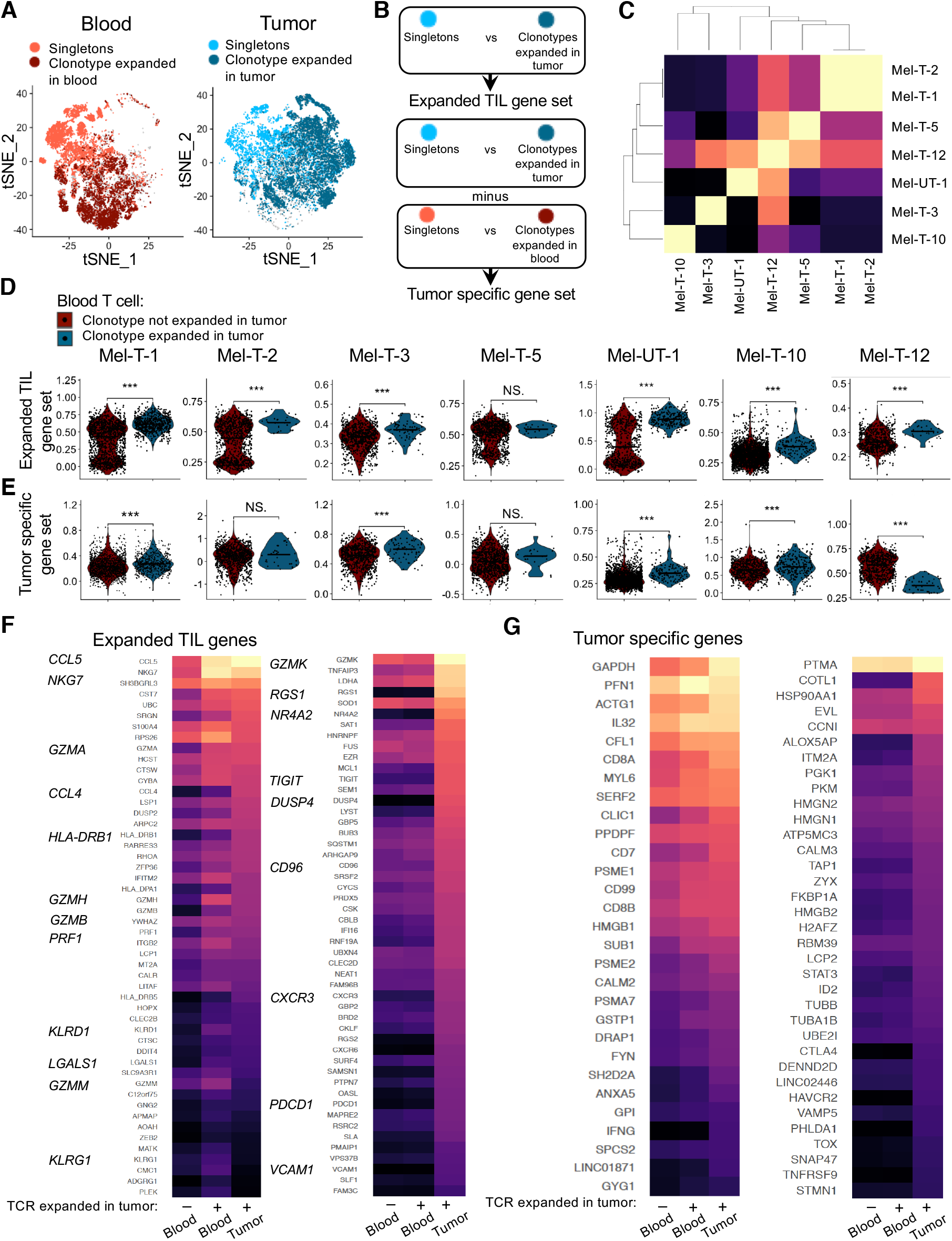
Circulating TILs reflect features of clonally expanded tumor-infiltrating T cells. **A**, tSNE plots highlighting tumor-infiltrating cells with a TCR clonally expanded in the tumor (dark blue) or singletons (light blue) and blood-derived cells with a TCR clonally expanded in blood (dark red) or singletons (tomato red). **B**, diagram illustrating the definition of expanded TIL gene sets and TIL-specific gene sets. **C**, clustered heatmap of the similarity matrix between expanded TIL gene sets of individual patients. **D**, quantification of the expression of expanded TIL gene sets in circulating TILs (blue) versus other circulating T cells (red) for each patient. Statistical significance was determined using Wilcoxon ranksum test. **E**, quantification of the expression of tumor specific gene sets in circulating TILs (blue) versus other circulating T cells (red) for each patient. Statistical significance was determined using the Wilcoxon rank-sum test. **F-G** for selected genes belonging the expanded TIL gene sets of at least three patients, expression is compared between tumor-resident clonally expanded TILs, circulating TILs and the rest of blood CD8 T cells. **F**, heatmap displaying expression of expanded TIL genes differentially expressed (left panel) or not (right panel) between circulating TILs and other circulating T cells. **G** heatmap displaying expression of tumor-specific genes differentially expressed (left panel) or not (right panel) between circulating TILs and other circulating T cells. For heatmaps, values are scaled and mean-centered log2-transformed gene counts. Statistical significance was determined with Wilcoxon rank-sum tests Bonferroni-corrected by the number of genes in the dataset (17.045) multiplied by the number of comparisons (3).

We then used expanded TIL gene sets to score individual CD8 T cells from the blood, and compared circulating TILs to other circulating T cells. Circulating TILs indeed had higher levels of aggregated expression of genes contained in expanded TIL gene sets in six out of seven patients (Fig 3D). Because expanded TIL gene sets contain multiple markers of activation, and, as shown in Fig 2A-B, circulating TILs are an effector population, it is possible that this result may just reflect a difference in activation. Therefore, we also measured the expression of tumor-specific gene sets in circulating TILs. For four out of seven patients, circulating TILs were also enriched for tumor-specific gene sets (Fig 3E). To visualize trends of expression of individual genes upregulated by clonally expanded TILs, we selected genes contained in the expanded TIL modules of at least three patients and compared differential expression between tumor-resident expanded TILs, circulating TILs and other circulating T cells. Of 566 genes tested, 283 were more highly expressed in circulating TILs compared to other circulating T cells, including multiple markers of effector functions. In contrast, 200 genes had similarly lower expression in blood compared to tumor regardless of TCR sharing. These genes included inhibitory receptors and other markers of terminal differentiation (Fig 3F, top 50 genes). Likewise, of 105 genes induced in the tumor-specific sets of at least two patients, 29 were significantly upregulated in circulating TILs, while 35 were not differentially expressed between blood populations. The tumor-specific genes reflected in circulating TILs included effector molecules (*IL32, IFNG*), enzymes involved in carbohydrate metabolism (*GAPDH, GPI, GYG1*) and genes involved in cytoskeleton dynamics (*ACTG, MYL6, PFN1, CFL1*), possibly mediating adaptations required for tissue residency (Kumar et al., 2017). Tumor-specific genes not reflected in circulating TILs also included some regulators of cell structure (*TUBA1B, TUBB, COTL1*), and additional inhibitory molecules (*HAVCR2, CTLA4*) and transcription factors associated with exhaustion (*TOX*) and tissue residency (*ID2*, Fig 3G). Thus, these data indicate that circulating TILs reflect certain features of the ongoing T cell response in the tumor including adaptations to tissue residency, but also reveal that key features of exhaustion in the tumor cannot be inferred from the periphery.

### Co-expression of KLRD1 and CD74 enriches for circulating TILs

We then examined whether circulating TILs can be identified in the blood based on expression of gene markers. We classified circulating CD8 T cells populations based on sharing of TCRs expanded in the tumor. The transcriptomes of individual cells with this annotation were used as the input of COMET (Combinatorial Marker Detection from Single-Cell Transcriptomic data, (Delaney et al., 2019), a tool that uses the minimal hypergeometric test to rank single and paired gene marker panels from a curated list of surface-expressed genes (Chihara et al., 2018) for their ability to identify predefined populations (Fig S3A). Of fourteen candidate gene marker pairs identified in five out of seven patients, the *KLRD1-CD74* module had the highest average rank and a highly significant q value in most patients (Fig S3B). Moreover, *KLRD1* and *CD74* were also highly ranked as candidate gene markers of circulating TILs in the single gene marker analysis for most patients (Fig S3C). We determined whether cells expressing high levels of the *KLRD1-CD74* module were enriched for circulating TILs. Indeed, selecting for cells that co-express *KLRD1* and *CD74* genes (Fig S3D) enriches for circulating TILs (Fig S3E). Interestingly, a significantly higher frequency of circulating TILs could not be found by classifying CD8 T cells by *PDCD1* gene expression alone (Fig S3F). Analysis of PD-1 expression at the protein level consistently revealed that while tumor-resident expanded TILs display higher levels of both *PDCD1* transcript and PD-1 protein than unexpanded cells, circulating TILs express similarly low levels of PD-1 as other circulating T cells (Fig S3G). Overall, these data suggest that it may be possible to identify a significant proportion of circulating T cells that are clonally related to tumor-resident T cells by measuring transcripts encoding for cell surface proteins in the absence of TCR sequence information.

### Tumor-blood sharing as a predictor of cytotoxic phenotype

Because many markers of exhaustion could not be detected in the circulation, we hypothesized that these are predominantly expressed by tumor-exclusive clones. To this end, we grouped CD8 TILs and circulating CD8 T cells into clonal groups, i.e. groups of cells sharing the same TCR, and calculated expansion and blood-tumor sharing scores for each clonal group. Mapping the tumor expansion and sharing scores onto the TIL tSNE plot shows clearly that most highly expanded clonal groups are not highly shared with the blood (Fig 4A-B).

**Fig. 4.**
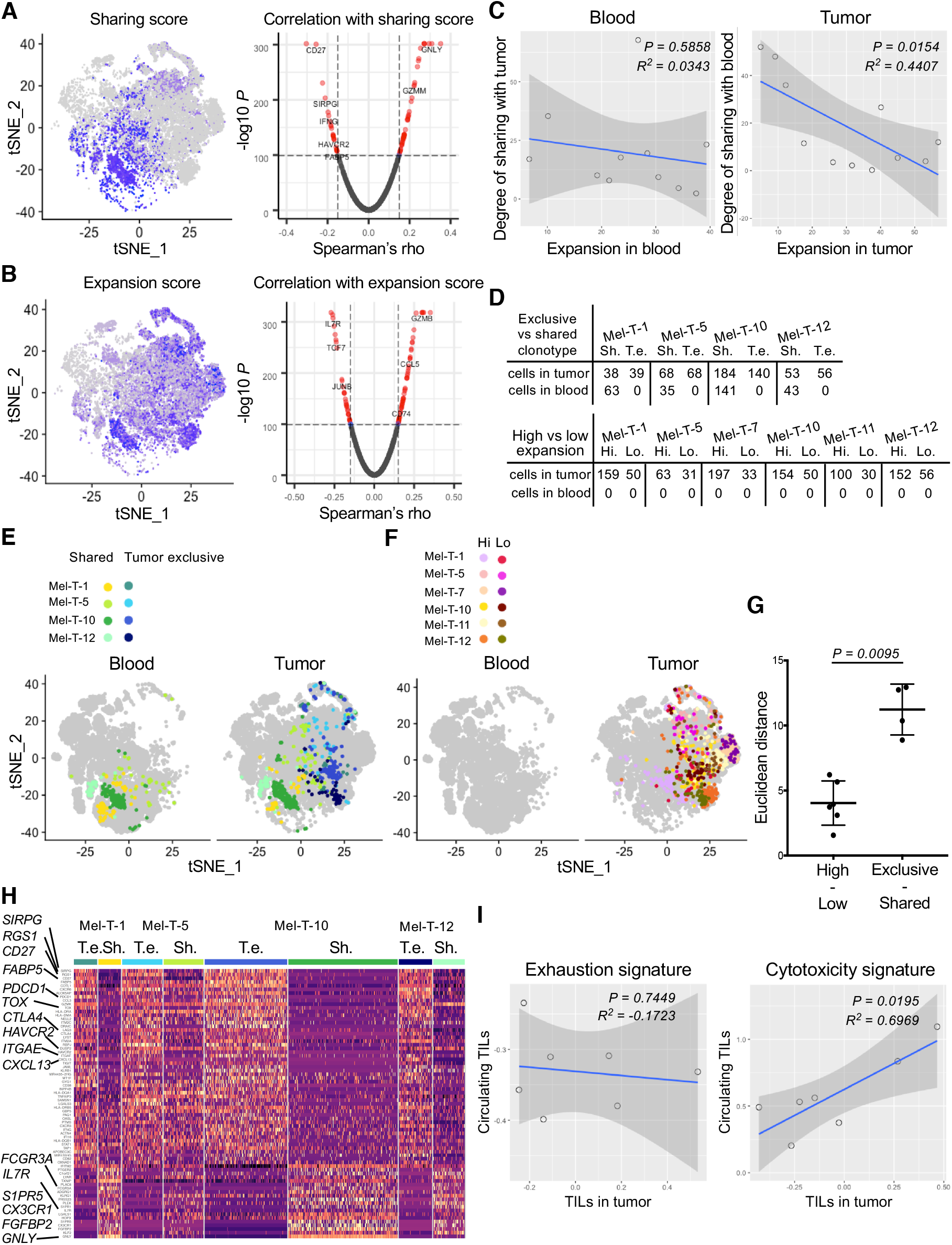
Tumor-blood sharing as a predictor of cytotoxic phenotype. **A**, tSNE plot of tumor CD8 T cells coloured by expression of a bood-tumor sharing score (left) calculated for each clonal group in the tumor (= cells sharing the same TCR) and volcano plot representing the Spearman’s rho coefficients and the adjusted p values of the correlations between sharing score and gene expression for CD8 TILs from 12 pooled patients (right). **B**, tSNE plot of tumor CD8 T cells coloured by expression of an expansion score (left) and volcano plot representing the Spearman’s rho coefficients and the adjusted p values of the correlations between expansion score and gene expression for CD8 TILs from 12 pooled patients (right). **C**, scatter plots showing the relationship between expansion score in the blood (left) or in the tumor (right) and degree of sharing across organ. Each dot represents the cumulative scores for each patient. Adjusted R-squared and F test p value are shown. **D**, table summarizing the number of cells expressing selected tumor exclusive (T. e.) and shared (Sh.) clonotypes in tumor and blood for the indicated patient (top). Table summarizing the number of cells expressing selected highly expanded (Hi) and lowly expanded (Lo) tumor-exclusive clonotypes in tumor and blood for the indicated patient (bottom). The indicated cells are mapped on a tSNE plot split by organ. **E**, **F** the indicated cells are mapped on a tSNE plot split by organ. **G**, Euclidean distances between the average of the PCA embeddings of each cell in the tumor exclusive versus shared clonal groups and highly expanded versus lowly expanded clonal groups. Distances were calculated using the PCs that were determined to be statistically significant in an elbow plot (8 PCs). Statistical significance between groups was determined using the Mann-Whitney test. **H**, heatmap displaying expression of genes differentially expressed between tumor exclusive versus shared clonal groups (rows, as in **E**). Cells are grouped and colour-coded according to the clonal group to which they belong. Differential expression was established with the Wilcoxon rank sum test, retaining genes with an adjusted p value < 0.05. Values are scaled and mean-centered log2-transformed gene counts. **I**, scatter plots showing the relationship of exhaustion signature (left) or cytotoxicity signature (right) between tumor infiltrating CD8 T cells and blood-derived circulating TILs. Adjusted R-squared and F test p value are shown.

We then determined whether gene expression in the tumor is associated with these metrics by calculating average Spearman’s rho coefficients for each gene. The expansion score appears to be correlated with an activation signature, with markers of naïve cells, (*IL7R, CCR7*, *LEF1*) displaying negative coefficients. Low values of the sharing score, instead, were associated with markers of exhaustion and terminal differentiation, (*HAVCR2, PDCD1, CXCL13, FABP5, IFNG*, Fig 4A-B). This approach did not allow us to separate the relative contribution of expansion and degree of sharing to these phenotypes. In fact, cumulative measures calculated for all the expanded clonal groups of each patient confirm that these two metrics are not independent, but rather inversely correlated, specifically in the tumor (Fig 4C).

To separate the effects of expansion and degree of sharing, we selected clonal groups for which either of these parameters was constant. Specifically, we identified four clonal group pairs with similar expansion scores within pairs, and opposite degrees of blood-tumor sharing. Similarly, we selected six clonal group pairs that were equally tumor exclusive and presented different degrees of clonal expansion (Fig 4D). We noted that pairs that differ by degree of sharing occupy distinct areas of the cell space, whereas pairs with variable expansion scores all map to the same area (Fig 4E-F). This difference could be quantified by calculating the average Euclidean distances for each clonal group pair. Indeed, the distances between exclusive versus shared clonal groups were significantly larger than the ones between highly versus lowly expanded clonal groups (Fig 4G). These data indicate that in the tumor infiltrate, the degree of sharing with blood explains more variation than the degree of clonal expansion. Moreover, we identified differentially expressed genes among cells belonging to these clonal groups and were able to confirm that TILs with a TCR shared with blood T cells were highly enriched for signatures of central memory and cytotoxicity, and presented less features of terminal exhaustion (Fig 4H).

As we demonstrated that circulating TILs express cytotoxic genes but low levels of exhaustion markers, we examined whether the cytotoxic score of circulating TILs was predictive of the cytotoxic score of the tumor infiltrate of the same patient. We scored individual tumor-derived CD8 TILs and circulating TILs for expression of a set of cytotoxicity markers previously reported in tumor-infiltrate T cells in single-cell RNAseq-based studies of multiple human cancers (see Methods and van der Leun et al., 2020). We observed a significant direct correlation between cytotoxicity scores in the tumor versus the circulating TIL population (Fig 4I). Moreover, this was not the case when scoring tumor-derived and circulating TILs for an exhaustion signature (Fig 4I), indicating that the degree of cytotoxicity of the tumor infiltrate, but not exhaustion, can be inferred from the periphery by analyzing circulating TILs.

### Signature of shared clones in the tissue of responders

It has been recently reported that clinical response to immunotherapy is accompanied by an influx of new T cell clones from the periphery into the tumor microenvironment, suggesting that TILs with a high degree of TCR sharing with blood can be involved in tumor rejection (Yost et al., 2019). To address this, we classified expanded TILs from the immunotherapy naïve patient Mel-T-1 into shared with blood or tumor exclusive, and scored individual cells for gene sets associated with response or disease progression after checkpoint immunotherapy (Sade-Feldman et al., 2018). While tumor exclusive TILs resembled TILs of progressors, genes predicting response were more represented in TILs shared with blood (Fig 5A-B). Furthermore, we examined whether defining sets of genes based on their correlation with the degree of blood-tumor sharing (Table S4) was sufficient to identify genes relevant to tumor rejection after immunotherapy. We analyzed residual tissue from two patients whose tumors responded clinically to checkpoint inhibitor immunotherapy. These samples did not contain any discernible tumor tissue, but rather focal aggregation of inflammatory cells including T cells and activated macrophages. We refer to these as tissue-infiltrating T cells. Similar to what we observed for tumor recurrent patients, in the two responder samples, less than 25% of tissue-infiltrating T cells carried a clonally expanded TCR (Fig 5C). We then analyzed the expression of genes that we had previously established to be associated with blood-tumor sharing or tumor-exclusivity in the tissue (Fig 4A, Table S5). Genes positively correlated with blood-tumor sharing were highly expressed in the clusters dominated by clonally expanded tissue-infiltrating T cells (Fig 5D-E). These included cytotoxic effector molecules and receptors (*GNLY*, *GZMH*, *GZMM*, *KLRD1*). Contrary to patients with recurrent disease, clonally expanded tissue-infiltrating T cells did not express high levels of co-inhibitory receptors (*TIGIT, HAVCR2, PDCD1, CTLA4*) and other markers of exhaustion (*TOX, TOX2, CXCL13*) that we demonstrated to be negatively correlated with the blood-tumor sharing score (Fig 5D-E). Overall, these results suggest that genes positively correlated with the degree of blood-tumor sharing mark T cells that are involved in tumor rejection in response to immunotherapy.

**Figure 5.**
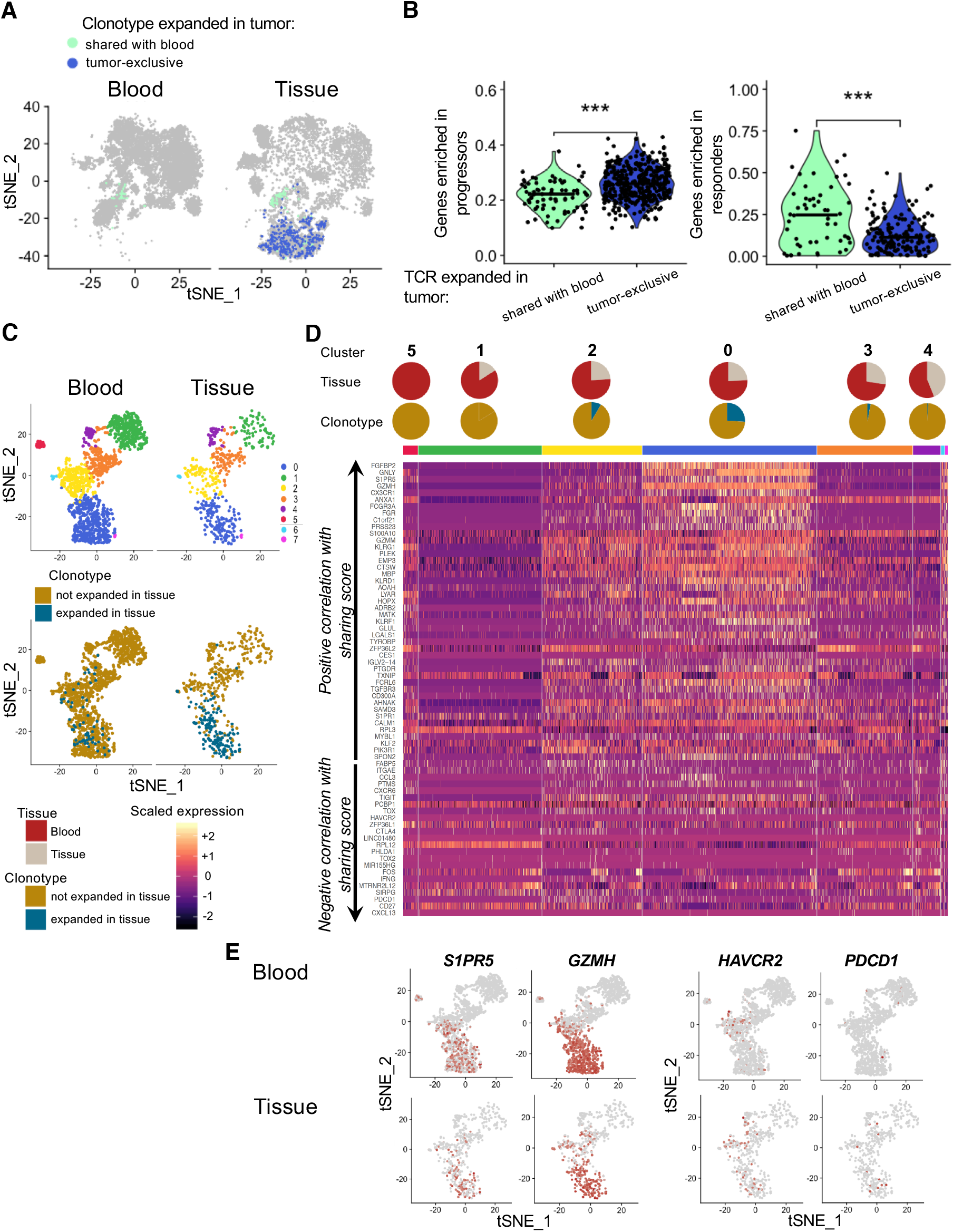
Features associated with blood-tumor sharing are highly expressed in the tissue of two patients who responded to immunotherapy. **A**, tSNE plots of blood and tumor T cells from patient Mel-UT-1 highlighting tumor-infiltrating cells with a TCR clonally expanded in the tumor and tumor-exclusive (dark blue) or shared with blood (aqua). **B**, quantification of the expression of gene sets associated with response to immunotherapy or progression as described by (Sade-Feldman et al. Cell 2019) in tumor-resident cells indicated in **A**. Statistical significance was determined using Wilcoxon rank-sum test. **C**, top, tSNE plot of CD8 T cells from 2 patients, split by tissue of origin and coloured by cluster. Clustering was obtained with a SNN modularity optimization-based algorithm (k parameter = 5, resolution = 0.5). Bottom, tSNE plot of CD8 T cells from 2 patients, split by tissue of origin and coloured by expression of a TCR expanded in tissue (blue), or not (gold). **D**, heatmap of genes previously established to be positively and negatively correlated with TCR sharing score at an adjusted Log(p value) <= −100 (as in Fig. 5A). Rows represent genes grouped by correlation with TCR sharing. Columns represent individual cells grouped by cluster identity (horizontal bars), with clusters ordered by decreasing percentage of blood cells. Values are scaled and mean-centered log2-transformed gene counts. Pie charts stacked on top of the grouping colour bar indicate tumor vs blood composition and expanded TIL composition for each cluster. E, tSNE plot of CD8 T cells from 2 patients, split by tissue of origin and coloured by scaled expression of the indicated genes.

Recent development in single-cell RNAseq technology has significantly improved our understanding of the relationship between TCR identity and functional properties. A number of studies have analyzed the overlap between TCR repertoires of discrete TIL clusters to examine developmental continuity across functional states (Tirosh et al., 2016; Li et al., 2019; Azizi et al., 2018). Here, we show that the degree of TCR overlap between the distinct tumor and blood tissue compartments can be indicative of the functional state of TILs. Specifically, we show that tumor-derived T cells belonging to TCR clonal groups highly shared across tumor and blood display the highest cytotoxic signatures in the tumor. Of note, circulating sister clones of these TILs also display cytotoxic functions proportionally to what is observed in the tumor, suggesting that analysis of circulating TILs could be used to predict effector functions in the tumor. Previous work using similar approaches in various cancers indicate that cytotoxicity represents one of two main developmental states of CD8 TILs, the other being T cell exhaustion (Zheng et al., 2017; Guo et al., 2018; Zhang et al., 2018; Azizi et al., 2018). These two phenotypes are conceptualized as the extremes in a bifurcation developmental model, where T cells belonging to the same clonal group tend to follow one or the other differentiation pathway. We consistently observe that cytotoxic and exhaustion phenotypes were also dichotomously associated with the pattern of TCR sharing across blood and tumor; while cytotoxic sister clones can be found in both tumor and blood, the most exhausted clonal groups appear to be tumor exclusive.

That the pattern of TCR sharing recapitulates CD8 T cell differentiation has a number of implications. In regard to the natural history of the antigen-specific response in the tumor, our data suggest that an exhausted phenotype is more likely to arise in T cells that have been primed locally in the tumor compared to central memory T cells seeded from the periphery. Indeed, tumor microenvironments contain antigen presenting cells that are unfit for priming T cell responses, partly as a consequence of interactions with regulatory T cells (Wolf et al., 2015; Magnuson et al., 2018).

The most exhausted TILs belong to tumor-exclusive clonal groups, implying that features of exhaustion cannot be inferred by analyzing circulating TILs. Nevertheless, it is known that tumor-resident exhausted T cells have only modest potential for functional reinvigoration, partly due to hard-wired epigenetic changes (Jadhav et al., 2019; Ghoneim et al., 2017; Pauken et al., 2016). Recent work has established that in metastatic melanoma patients, clinical response to checkpoint inhibition immunotherapy is associated with peripheral expansion of large clonal groups expressing high levels of cytotoxic and effector markers, with strong overlap with genes that we found to be associated with a high degree of blood-tumor sharing (Fairfax et al., 2020). This observation would suggest that circulating TILs contribute to response to immunotherapy, although the specificity of this population for tumor antigens remains to be demonstrated. Moreover, a study of basal cell carcinoma lesions sampled before and after anti-PD1 therapy found that pre-existing exhausted TILs were largely replaced by novel expanded clonotypes, reinforcing the idea that exhausted clonal groups might be short-lived and unlikely to participate in long-term immune surveillance (Yost et al., 2019). We report that gene associated with high degree of blood-tumor sharing are enriched for effector signatures. Consistently, a recent meta-analysis of four clinical trials of an anti-PD-L1 antibody (atezolizumab) in a variety of solid tumors found that in the tumor, gene signatures associated with clonotypes that were expanded in both tumor and normal adjacent tissue predicted a better response to anti-PD-L1 (Wu et al., 2020).

As we wished to examine the circulating TILs in a variety of clinical settings, we did not select patients by prior exposure to immunotherapy or variability in clinical presentation. We thus were able to focus on findings that were reproducible in the patients analyzed, with the goal that these findings could be further corroborated in larger, uniformly treated patient cohorts. While we have further shown that it could be possible to enrich for circulating TILs by analyzing co-expression of genes encoding for surface markers, critical validation steps are required for implementation into a scalable immune monitoring assay. Nevertheless, we note that, in a companion paper by Pauken et al., circulating TILs could be identified by the expression of a set of gene markers partially overlapping with the ones that we found. The authors describe a combination of four negation markers (*CCR7*, *FLT3LG*, *LTB*, and *GYPC*) that allows for enrichment of circulating TILs in treatment naïve metastatic melanoma patients. Indeed, three of these markers were also highly significantly associated with circulating TILs in our dataset in the two patients with the highest frequency of circulating TILs. Therefore, it appears that the approach of using the TCR as a molecular barcode for circulating TILs followed by an unbiased search for candidate markers can return the same genes across distinct patient cohorts and independent laboratories. Finally, we could not assess the directionality of circulating TILs with respect to their tumor-resident sister clones to determine whether circulating TILs include cells that have left the tumor bed and re-entered the circulation. This possibility has been described for tissue-resident memory T cells in the skin: a subset of these cells can enter the circulation, where they remain transcriptionally representative of the skin-resident population (Klicznik et al., 2019). Additional work is required to assess if a similar phenomenon also occurs in cancer, and whether related clonal T cells can sequentially patrol multiple metastases, including tissues to which immune cell trafficking is tightly regulated, such as the central nervous system.

In conclusion, we propose that in human cancer the pattern of TCR repertoire sharing across tissues recapitulates T cell differentiation states. Moreover, we report that it is possible to assess the ongoing immune response in the tumor by analyzing a circulating T cell population. Further work is necessary to better define circulating TILs as blood-based biomarkers to increase the precision of existing ‘liquid biopsy’ assays.

## Author contributions

L.E.L. planned and performed experiments, analyzed data and wrote the manuscript with inputs from P-P. A., B.L., B.H., J.S., M.S., H.K. and D.A.H; P-P. A. and B.L. analyzed data; B.H. performed experiments, L.Z. and K.R. provided technical help with single-cell genomics; L.Z. provided technical help with sample acquisition; K.O. and J.C. harvested patient tissue samples, S.J. collected and reviewed patient clinical information; M.S. oversaw data analysis; L.E.L., M.S., H.K. and D.A.H. conceptualized the study; H.K. and D.A.H. planned and supervised the study and acquired funding.

## Acknowledgements

The authors would like to thank Lesley Devine and Chao Wang for technical assistance with flow cytometry, Guilin Wang and the staff of the Yale Center for Genome Analysis for technical assistance with single cell genomics, and Antonella Bacchiocchi from the Yale SPORE in Skin Cancer Biospecimen Core for assistance with sample collection. The authors would also like to thank Philip Coish and Jessica Wei for help with manuscript preparation and Matthew Lincoln and Rahul Dhodapkar for insightful comments on data analysis.

## Funding Sources

This work was supported by NIH grant P50 CA121974 (to Harriet Kluger and Marcus Bosenberg). and NIH grants P01 AI073748, U24 AI11867, R01 AI22220, UM 1HG009390, P01 AI039671, P50 CA121974, R01 CA227473) to DAH.

## Conflicts of interest

Though we believe that none of these relationships are conflicts of interest, Dr. Hafler has received research funding from Bristol-Myers Squibb, Sanofi, and Genentech for work unrelated to this project. He has been a consultant over the past ten years for Bristol Myers Squibb, Compass Therapeutics, EMD Serono, Genentech, Juno therapeutics, Novartis Pharmaceuticals, Proclara Biosciences, Sage Therapeutics, and Sanofi Genzyme. Dr. Kluger has received research funding from Bristol-Myers Squibb, Apexigen and Merck. She has been a consultant for Corvus, Biodesix, Roche-Genentech, Pfizer, Iovance, Immunocore, Celldex, Array Biopharma, Merck, Elevate Bio and Instil Bio.

## Materials and Methods

### Human subjects

This study was approved by the Yale University Institutional Review Boards. All participants provided written informed consent for the study. All patients had histologically confirmed Stage IV metastatic melanoma. Surgical resection was clinically indicated in all cases. Patients donated a portion of surgically resected tissue (for TIL isolation) and a time-matched blood sample (for PBMC isolation).

### Tumor dissociation and PBMC isolation

Freshly resected tumor samples were mechanically dissociated and digested in HBSS medium with Collagenase IV (2.5 mg/mL) and DNAse I (0.2 mg/mL) (Worthington Biochemical Corporation) at 37°C for 30 minutes. TIL-enriched single cell suspensions were then isolated using Lymphoprep gradient centrifugation. PBMCs were isolated from whole blood using Lymphoprep gradient centrifugation. For sample Mel-T-9, TILs and PBMCs were cryopreserved in GemCell human AB serum with 10% DMSO in liquid nitrogen. For sample Mel-T-11, cryopreserved TILs and PBMCs were kindly provided by the Advanced Cell Therapy Laboratory in the Department of Laboratory Medicine (Yale School of Medicine).

### Cell sorting

All samples but Mel-T-9 and Mel-T-11 were sorted directly after PBMC and TIL isolation. For sample Mel-T-9 and Mel-T-11, cryopreserved TIL and PBMC aliquots were thawed following the protocol CG00039 from 10X Genomics (https://support.10xgenomics.com/). Samples were sorted on a BD FACS Aria II using a live cell (Live/Dead Cell Viability Assay, Life Technologies), CD45+ (eBioscience, PerCP-Cy5.5 conjugated), TCRαβ+(eBioscience, PE-Cy7 conjugated) gating strategy. At least 5,000 live T cells were sorted per sample for inclusion in the 10X library preparation.

### Preparation of scRNAseq and scTCRseq libraries

Sorted T cell samples were centrifuged at 300 rcf for 10 minutes at 4°C, resuspended in PBS with 0.04% BSA and counted on a hemocytometer. Cell concentrations were adjusted to be between 700 and 1,000 cells/μL. Paired single-cell gene expression and TCR libraries were prepared and sequenced at the Yale Center for Genome Analysis following standard protocols from 10X genomics (https://medicine.yale.edu/keck/ycga/sequencing/10x/singcellsequencing/). Briefly, single cells were isolated in 1 nanoliter GEMs (Gel Bead in Emulsion), using the GemCode technology. Depending on the initial cellularity of the sorted samples, between 5,000 and 10,000 cells were targeted. Cell barcoding, lysis, and reverse transcription of mRNA occurred within each GEM. cDNA libraries were then generated using next-generation sequencing PCR amplification with the 5-prime chemistry. Both scRNAseq libraries and scTCRseq libraries were generated for each sample.

### CITE-seq staining

Sorted live/CD45+/TCRαβ+ cells derived from blood and tumor samples were incubated with 0.5 μg/sample of the following antibody mix: αCD4 clone RPA-T4, αCD8a clone RPA-T8, aPD-1 clone EH12.2H7 (all from BioLegend, conjugated with TotalSeq™ oligos) in PBS:BSA 0.1% for 30 minutes on ice. Stained cells were centrifuged at 300 rcf for 10 minutes at 4°C, resuspended in PBS with 0.04% BSA, counted on a hemocytometer and used for preparation of paired scRNA-seq, scTCR-seq and scCITE-seq libraries, as described above.

### Sequencing

scRNAseq libraries were sequenced on an Illumina NovaSeq S4 instrument at a read length of 26×8×91 base pairs and at a depth of 300 million reads per sample. CITE-seq libraries were also sequenced on at a read length of 26×8×91 base pairs and at a depth of 30 million reads per sample scTCRseq libraries were sequenced on an Illumina NovaSeq S4 at a read length of 150×150 base pairs and at a depth of 20 million reads per sample.

### Data processing of scRNAseq and scTCRseq libraries

ScRNAseq reads were aligned to the reference genome GRCh38 using the cellranger count pipeline from the CellRanger software v3.0.2 (10X Genomics). The mean number of reads per cell was 49,499 (28,471 – 103,776) for blood T cell samples, and 92,357 (43,243 – 283,598) for tumor T cell samples. The median number of unique molecular identifiers (UMI) per cell was 4,383 (2,624 – 5,820) for blood and 3,960 (2,440 – 4,836) for tumor samples. The median number of unique genes per cell was 1,444 (1,105 – 1,802) for blood and 1,465 (1,060 – 1791) for tumor samples. Filtered gene-barcode matrices containing barcodes that had passed the threshold for cell detection were used for downstream analyses.

ScTCRseq reads were processed using the cellranger vdj pipeline from CellRanger v3.0.2 (10X Genomics, reference genome GRCh38). The mean number of read pairs per cell was 4,792 (1,358 – 10,040) for blood-derived TCR libraries and 6,442 (1,765 – 27,411) for tumor-derived TCR-libraries. The VDJ enrichment, defined as the percentage of reads mapping to any VDJ gene, was 89% ± 2.4 for blood and 90% ± 2.58 for tumor. TCR contigs called by cellranger vdj with high confidence, listed in the filtered_contig_annotations.csv output file, were retained for downstream analyses. Nucleotide and amino acid CDR3α and β sequences and count matrices of gene expression were matched for each cell based on barcode identities using costume R scripts and imported into Seurat v3.0 (www.satijalab.org/seurat/). When more than one CDR3α and β sequence was found for a given cell, additional sequences were appended, while NA values were assigned for missing sequences. For downstream analyses, all cells for which multiple CDR3α and β sequences existed or NA values were present were removed from the dataset.

### Cluster analysis

Cell-level quality control was performed for individual patients and for blood and tumor gene expression matrices separately. Thresholds for removing low-quality cells were selected based on the distribution of the number of unique genes per cell and the percentage of genes that map to the mitochondrial genome. On average, cells that expressed between 258 and 3,896 unique genes and with a percentage of mitochondrial reads below 13.75% were retained for downstream analyses.

For the cluster analysis, we filtered out T cell receptor genes and genes encoded by the Y chromosome and selected for cells with *CD8A* count >= 1 and *CD3E* + *CD3D* + *CD3G* >= 1. We then joined count matrices from individual patients to generate one merged Seurat object for blood-derived cells and one for tumor-derived cells. Counts were log-normalized and 2,000 variable genes were identified for blood and tumor separately using the Seurat function FindVariableFeatures with the “vst” method (variance stabilizing transformation). Tumor and blood datasets were subsequently integrated into a single dataset using the Seurat functions FindlntegrationAnchors and IntegrateData using 20 dimensions. For the resulting integrated object, scaled z-scores for each gene were computed using the ScaleData function and regressed against sequencing batch and fresh or frozen processing. Principal component analysis was then run to identify 30 principal components, of which the first 8 were selected for t-SNE dimensionality reduction using the elbow method on the distribution of the standard deviation of each PC. For clusters identification, we initially used the FindNeighbors function to build a shared nearest neighbor graph based on the k-nearest neighbor of each cell, with k = 5. Clusters are then determined with the FindClusters function, which uses the Louvain algorithm for modularity optimization (ref, resolution = 0.4). Markers of individual clusters were defined as the differentially expressed genes between cells belonging to each cluster versus every other cell using the Wilcoxon Rank Sum test as implemented in the function FindMarkers. Genes with an adjusted p value < 0.05 were retained.

### TCRαβ repertoire analysis

The TCRαβ repertoires of blood and tumor samples were analyzed using scRepertoire (Borcherding et al., 2020). Filtered contig annotation files from the output of cellranger vdj were used as input. Indexes of diversity (Shannon entropy, Inverted Simpson) and species richness (Chao, ACE) were calculated using the clonalDiversity function (cloneCall = “aa”) and the significance of group differences was tested with the Wilcoxon rank-sum test. Hierarchical clustering of samples based on clone size distribution was performed with the function clonesizeDistribution (cloneCall = “aa”) derived from the package PowerTCR (Koch et al., 2018). The proportion of occupied repertoire space was plotted using the function clonalProportion (cloneCall = “aa”) and alluvial plots were generated with compareClonotypes after selecting the top 20 most expanded clonotypes in the tumor.

To obtain a binary definition of clonal expansion, thresholds were selected by inspection of histograms displaying the percentage of the tumor TCRαβ repertoire occupied by a given clonotype on the y axis, and individual TIL clonotypes ordered by increasing percentage of the tumor TIL repertoire on the x axis to identify a knee of the curve.

### TIL expanded gene set analysis

Gene sets of clonally expanded TILs were calculated for each individual patient. To determine which genes are specifically associated with T cell expansion in the tumor we modeled raw gene counts in a generalized linear model as implemented in DESeq2 (Love et al., 2014) including an interaction term between the expansion and the tissue variable: ~Tissue+Expansion+Tissue:Expansion. We ran our model on T cells from each patient separately, using a local fit. To generate expanded TIL gene sets, we filtered genes with log2FoldChange_tumor > 0 at an adjusted p value < 0.1. To generate tumor-specific gene sets, we filtered genes with log2FoldChange_tumor > 0 at an adjusted p value < 0.1 and log2FoldChange_interaction >0 at an adjusted p value <0.1. We then used the function AddModuleScore with default parameters to calculate the average expression of each gene set on single cells from the blood, for each patient. Blood cells were annotated based on TCR identity as circulating TILs (those with a TCR found to be above the threshold for clonal expansion in the tumor) and or not. Statistical significance of the difference between the means of each group of cells was established for each patient using the Wilcoxon rank-sum test.

### COMET analysis

The input of the COMET analysis consisted of log-normalized counts of CD8 T cells from the blood of individual patients. Each cell was annotated as having or not a TCR clonally expanded in the tumor. Expression matrices and cell identity annotations were uploaded to http://www.cometsc.com/comet and comet was run with default parameters. For each patient, we obtained an output file containing lists of gene pairs ranked by their power to identify circulating TILs. We joined these lists by the gene pair identity and filtered out all gene pairs that were not detected in at least 5 out of 7 patients. For the remaining gene pairs, we calculated an average rank and ordered them by decreasing values of the average rank, the highest of which corresponded to the gene pair *KLRD1-CD74*.

### Expansion and blood-tumor sharing score

At the clonotype level, the expansion score (Fig 4B) was defined as follows. For each clonotype, we calculated the percentage of the total tumor T cell infiltrate occupied by cells sharing that same clonotype (clonal groups). The patient-level expansion score, as displayed in Fig 4C, was calculated by adding up the expansion scores of individual clonal groups that were identified as clonally expanded as exemplified in Fig 1A. Separate patient-level expansion scores were calculated for blood and tumor. The sharing score of individual tumor-expanded TCRs (Fig 4A) was defined as the percentage of cells in each clonal group that were also found in blood. For blood-expanded TCRs, the sharing score was a measure of the frequency of cells with that TCRs that were also found in the tumor infiltrate. At the patient-level, the degree of sharing was calculated by averaging the sharing score of individual clonal groups that were identified as clonally expanded as exemplified in Fig 1A. Separate patient-level sharing scores were calculated for blood and tumor.

### Exhaustion and cytotoxicity gene signatures

Consensus markers of exhaustion and cytotoxicity were obtained from a recent review of CD8 states in human cancer (van der Leun et al., 2020). Specifically, the exhaustion gene set included *HAVCR2, IFNG, ITGAE, PDCD1, CXCL13, LAYN, LAG3, TIGIT, CTLA4, ENTPD1*, while the cytotoxicity gene set consisted of *CX3CR1*, *PRF1*, *GZMA*, *GZMH*, *GNLY*, *FGFBP2*, *KLRG1*, *GZMM, LYAR, TXNIP, FCRL6, NKG7, KLRD1*. We then used the function AddModuleScore with default parameters to calculate the average expression of each gene set on single cells. For each patient, we averaged the exhaustion and cytotoxicity scores of all tumor-infiltrating CD8 T cells and circulating TILs.

## Supplementary Materials

Fig. S1 Individual thresholds of clonal expansion in the tumor.

Fig. S2 Individual thresholds of clonal expansion in the blood.

Fig. S3 Co-expression of KLRD1 and CD74 enriches for circulating TILs.

Table S1 Top 10 marker genes for each cluster, with percentage of positive cells in the cluster of interest, delta percentage of positive cells between that cluster and the rest of the dataset, and adjusted p-value.

Table S2 List of gene marker pairs of circulating TILs found in at least five out of seven patients, with rank and q value for each patient.

Table S3 Expanded TIL gene sets and tumor-specific gene sets for each patient.

Table S4 List of correlation coefficients between genes and expansion score, filtered for −LogP >= 100. Same for sharing score.

Table S5 Amino acid and nucleotide sequences of the TCRαβ CDR3 regions of clonotypes displayed in Figure 4.

---

**Fig. S1 Individual thresholds of clonal expansion in the tumor.** Bar graphs displaying the frequency of TILs expressing a given TCRαβ clonotype on the y axis, and individual TIL clonotypes ordered by increasing frequency on the x axis, for individual patients.

**Fig. S2 Individual thresholds of clonal expansion in blood.** Bar graphs displaying the frequency of TILs expressing a given TCRαβ clonotype on the y axis, and individual TIL clonotypes ordered by increasing frequency on the x axis, for individual patients.

**Fig. S3 Co-expression of KLRD1 and CD74 enriches for circulating TILs. A,** workflow for identification of gene markers of circulating TILs. Singleton and paired gene markers of circulating TILs were identified from the blood of each patient using COMET. Gene marker pairs detected in most patients were chosen and blood T cells were classified based on co-expression of the selected gene markers. The percentage of circulating TILs in cells positive and negative for the selected gene markers was compared for each patient. **B**, table summarizing the rank of the gene marker pair *KLRD1-CD74*, the q value and the total number of gene marker pairs identified by the COMET analysis. N.D. indicates that the gene marker pair *KLRD1-CD74* was not detected in that patient. N.A. = not applicable. **C**, heatmap of the reverse rank (rank * −1, rows) of singleton gene markers of tumor directed T cells that were shared by at least 5 out of 7 patients (columns), ordered by the average across patients. For pairs not detected in a specific patient, a reverse rank of −200 was assigned. Arrows indicate *KLRD1* and *CD74*, detected by both the singleton and paired gene marker analysis. **D**, scatter plots displaying the relationship between the integrated (Stuart et al., 2019) expression of *KLRD1* and *CD74* transcripts in each patient. Dashed boxes indicate double positive cells, with expression of *KLRD1* >= 1.5 and *CD74* >= 3. **E**, percentage of cells expressing a TCR expanded in the tumor among cells that are positive or negative for the *KLRD1-CD74* combination in each patent. **F**, percentage of cells expressing a TCR expanded in the tumor among cells that are positive or negative for *PDCD1* expression in each patent. Statistical significance was determined using paired t tests. **G**, Pie charts displaying the percentage composition in terms of TCRs expanded in the tumor, shared but not expanded in the tumor, and not shared with the tumor for blood T cells classified as positive or negative for the *KLRD1-CD74* marker combination. Percentages are calculated on the sum of all blood CD8 T cells from the 7 patients.

