## Supplementary Figures 1-3 for "Circulating Clonally Expanded T Cells Reflect Functions of Tumor Infiltrating T Cells"

**Figure S1**

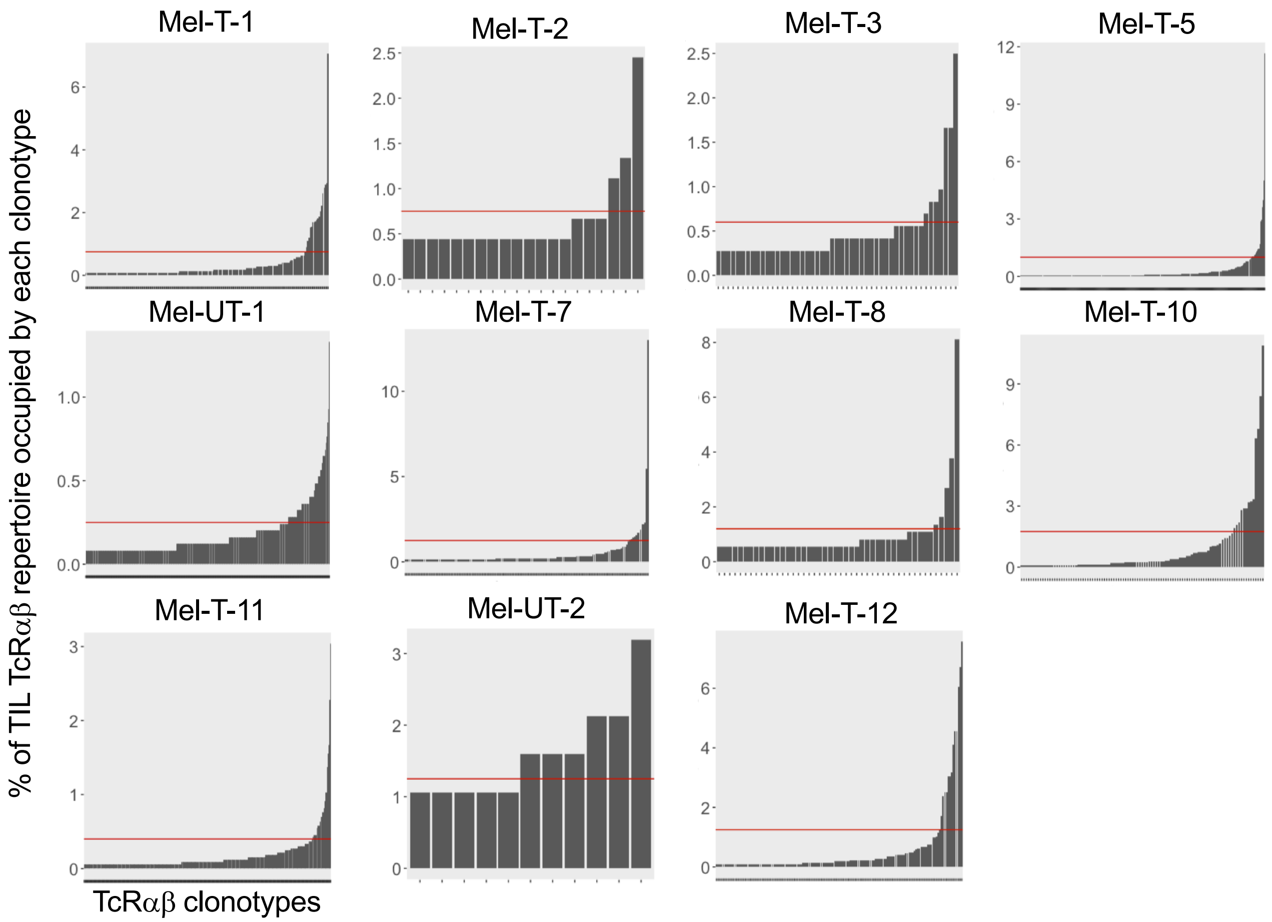

Mel-T-1
Mel-T-2
Mel-T-3
Mel-T-5
Mel-UT-1
Mel-T-7
Mel-T-8
Mel-T-10
Mel-T-11
Mel-UT-2
Mel-T-12
% of TIL TcRαβ repertoire occupied by each clonotype
TcRαβ clonotypes

**Figure S2**

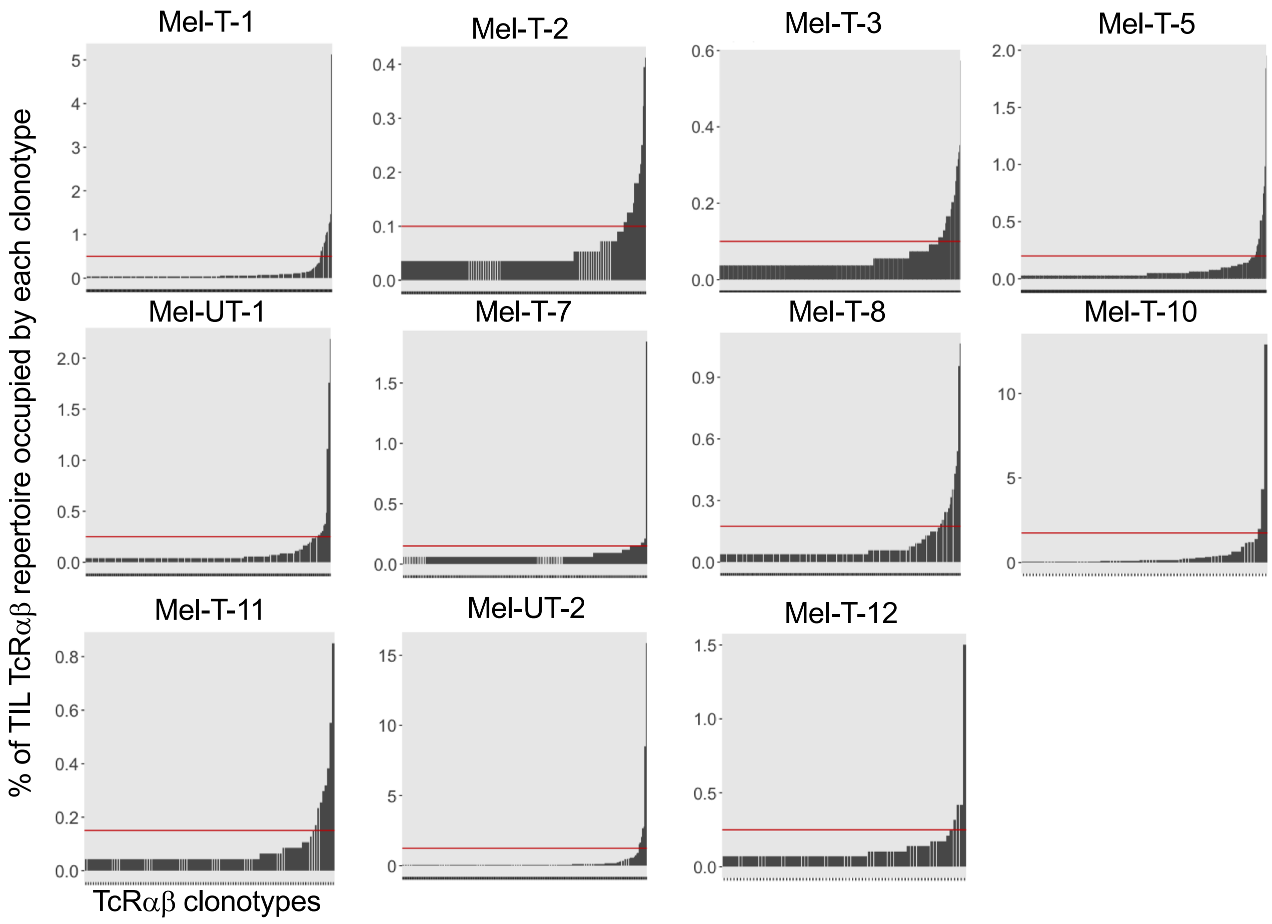

Mel-T-1
Mel-T-2
Mel-T-3
Mel-T-5
Mel-UT-1
Mel-T-7
Mel-T-8
Mel-T-10
Mel-T-11
Mel-UT-2
Mel-T-12
% of TIL TcRαβ repertoire occupied by each clonotype
TcRαβ clonotypes

### Figure S3

A

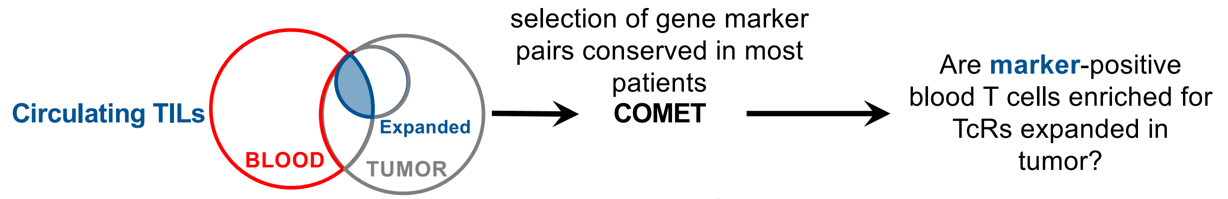


B

Paired gene markers of **circulating TILs**

| ***CD74 – KLRD1*** | Mel-T-1 | Mel-T-2 | Mel-T-3 | Mel-T-5 | Mel-UT-1 | Mel-T-10 | Mel-T-12 |
|---|---|---|---|---|---|---|---|
| Rank (Total # of pairs) | 305 (2000) | 33 (91) | 63 (2000) | 12 (1485) | N.D. (6) | 578 (2000) | N.D. (2000) |
| Q value | $4.31*e^{-40}$ | 0.289 | 0.0003 | $2.73*e^{-5}$ | N.A. | $1.78*e^{-14}$ | N.A. |

C

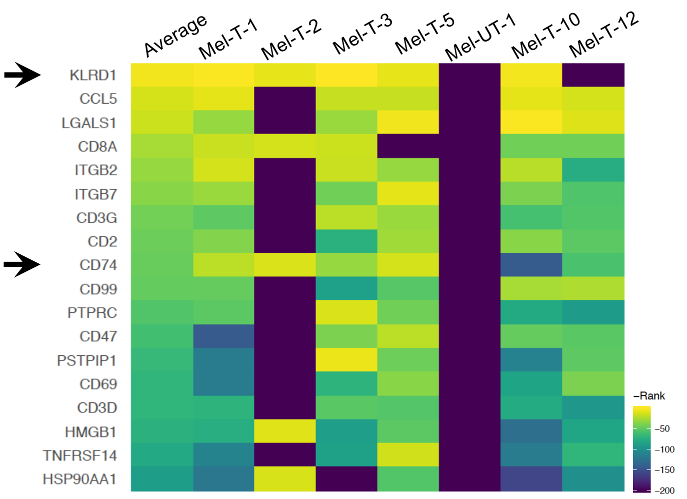


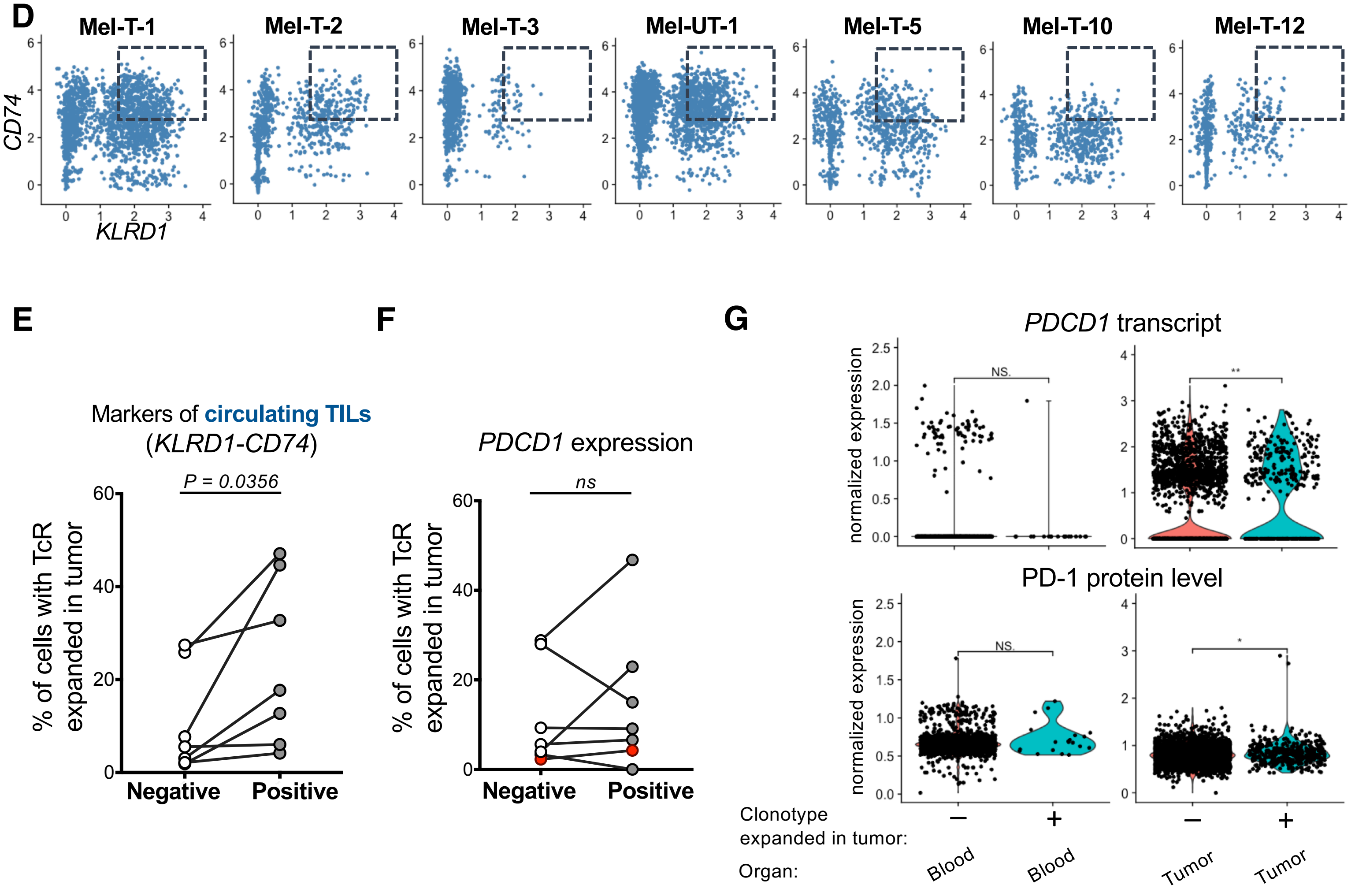
